# A functional pre-screening platform for identifying points of vulnerability in the cell death map of human melanoma tumors

**DOI:** 10.1101/2020.11.11.377671

**Authors:** Naama Pnina Dekel-Bird, Shani Bialik, Orit Itzhaki, Tomer Meir Salame, Naama Yaeli-Slonim, Vered Levin-Salomon, Santosh Kumar Dasari, Michal Besser, Adi Kimchi

## Abstract

Targeted drug therapy in melanoma patients carrying the *BRAF* V600E mutation provides temporary remission, often followed by relapse due to acquired drug resistance. Here we propose a functional approach to circumvent drug resistance by applying a personalized prescreening platform that maps points of vulnerability in each tumor, prior to drug treatment. This platform applies siRNAs targeting 81 apoptosis, autophagy and programmed necrosis genes in patient tumor cell cultures, identifying genes whose targeting maximizes cell killing by short-term BRAF inhibition. Melanoma tumors displayed large heterogeneity in the number and identities of soft-spots, providing different tumor-specific functional death signatures. The soft-spots were targeted by replacing functional siRNAs with small compound inhibitors for long-term treatment in combination with vemurafenib. This strategy reduced the number of drug-tolerant persister cells surviving treatment, and most importantly, the number of drug-resistant foci. Thus, prescreening melanoma tumors for soft-spots within the cell death network may enhance targeted drug therapy before resistance emerges, thereby reducing the odds of developing drug-resistant mutations, and preventing tumor relapse.

## Introduction

Immunotherapy and targeted therapy are the two main therapeutic arms currently available for treating melanoma. While over recent years, immune checkpoint inhibitors have been a breakthrough in the treatment of patients with advanced melanoma, a substantial portion of patients are refractory to or progress after treatment (1,2). A major difficulty also persists for targeted therapy, which is based on drugs targeting the mitogen-activated protein kinase (MAPK) signaling pathway that dominates melanoma tumors. In 50% and 20% of melanoma patients, the MAPK pathway is stimulated by mutations in *BRAF* or *NRAS*, respectively (3). Specific drugs targeting the most abundant *BRAF* V600E mutation (e.g., vemurafenib (4)), MEK (MAP2K) inhibitors (5) and ERK1/2 (MAPK1/3) inhibitors (6) have been developed and used in the clinic. While very effective during the first months of treatment, in a significant fraction of patients, positive tumor responses are followed by a relapse due to the development of acquired drug resistance. Combining MEK inhibitors with BRAF inhibitors from the start of treatment delays the time until relapse, but does not prevent it (7). While overall, current treatment strategies have improved 5-year survival rates of patients with metastatic melanoma from 5-10% to around 50%, there is still great need for improving the efficacy of standard target therapy to raise survival rates.

Acquired drug resistance develops from persister cells, sub-populations of cancer cells that maintain viability under continuous drug treatment (8–12). These cells do not harbor inheritable genetic or stable epigenetic modifications, and their drug tolerance phenotype is reversible. They may replicate at low frequency or remain quiescent in the continuous presence of drug. The persister cells are a dangerous cellular reservoir from which acquired mutations that bypass the effect of the drug and confer drug resistance develop during prolonged treatments. Much effort has been expended in recent years in identifying these mutations, and in designing accordingly a second front of drug treatment that targets the new mutations. For example, acquired resistance to BRAF inhibitors has been attributed to activating mutations in *MEK1* (13) or *MEK2* (14), mutated *BRAF* splice variants with increased kinase activity (15) and overexpression of mutated BRAF itself. Unfortunately, at this stage, tumors are increasingly refractory even to targeted treatment. Yet due to the multiplicity of such mutations, and their consequent unpredictable nature, mechanisms of potential resistance cannot be targeted at earlier stages.

Here we suggest an alternative way to address the problem of acquired drug resistance by identifying points of vulnerability in melanoma tumors right at the start of vemurafenib treatment, the targeting of which would reduce the number of persisters and as a consequence, the odds of developing drug resistant mutations. It is based on the premise that the suppression of the MEK/ERK pathway by BRAF inhibitors can be wired in different ways to various molecular mechanisms that ultimately reduce tumor cell number, i.e. different pathways that control the life and death decisions of a cell, such as apoptosis, necrosis, and autophagy (16–20). Our challenge was to identify the proteins functioning along these pathways, whose reduced expression or function increases the killing effects of the targeting drug (i.e., the soft-spots in these pathways). As multiple interconnected pathways control this important balance (21), and each patient’s tumor may carry different genetic/epigenetic alterations that affect the functionality of some of these pathways, we performed a personalized functional search for identifying these points of vulnerability. The screen covered 81 genes operating along apoptosis, autophagy and programmed necrosis pathways, and measured the outcome of their knock-down, one at a time, after 24h of drug treatment in early passage melanoma cell cultures carrying the *BRAF* V600E mutation, prepared from a cohort of 12 BRAF inhibitor naïve patients. Using this platform, we identified a large heterogeneity among individual tumors, differing in the number and the identity of points of vulnerability upon vemurafenib treatment. Targeting the soft-spots identified in this manner led to significant reductions in the number of drug tolerant cells that maintained their survival during long term drug exposure, and most importantly, attenuated the subsequent development of drug resistant foci that appear after several weeks of drug treatment.

## Materials and Methods

### Reagents, cell culture and transfections

All chemical reagents were purchased from Sigma-Aldrich unless otherwise stated. Primary melanoma lines were generated from tumor biopsies of metastatic melanoma patients, as detailed in Supplementary Table S1, enrolled in a phase II TIL ACT trial at the Sheba Medical Center. Patients signed an informed consent approved by the Israeli Ministry of Health (Helsinki approval no. 3518/2004, NCT00287131), which allowed enrollment in the clinical trial and the use of excess cell material for research purpose. Fragmentation, enzymatic digestion and tissue remnant culture techniques were used to isolate melanoma cells from surgically resected metastatic lesions to generate primary melanoma cell lines (22). These cell lines, which tested negative for mycoplasma at thawing, were maintained in T2 medium (RPMI, 10% FBS (HyClone, Thermo Scientific, SH3010903), 1mM sodium pyruvate, 25mM HEPES, Pen/Strep, L-Glutamine (all from Biological Industries). The cells were treated with vemurafenib at the indicated concentrations (Selleckchem, S1267), 0.5μM or 1μM S63845 (ApexBio, A8737-5), 0.5μM or 1μM ABT-737 (Santa Cruz, sc-207242), 20μM HCQ (Sigma, C6628, H0915), 5-10nM bafilomycin A1 (LC-Laboratories, B-1080), 0.5μM or 1μM venetoclax (Medchemexpress, HY-15531) and 0.5μM or 1μM A-1331852 (Medchemexpress, HY-19741). For transient transfections of siRNA, 25nM siGNOME siRNA pool (Dharmacon), or a total of 50nM for double KD, was mixed with transfection reagent Dharmafect1 (Dharmacon) and added to the cells 24h post plating.

### Cell viability and death assays

Cell viability was measured using luminescence based CellTiter-Glo assay according to the manufacturer’s instructions (Promega, G7572). Luminescence was read in a microplate luminometer (TECAN, Tecan Trading AG). For calcein/PI assays, 2.5×10^5^ cells were grown in 6-well plates and 24h post treatment, 1μM Calcein AM (Life Technologies #C3099) and 1.5μM PI (Sigma, P4864) were added. The plates were then incubated for 30 min at 37°C and live and dead cells were viewed by fluorescence microscopy (Olympus BX41 or Olympus IX73 microscopes) equipped with 10x or 20x objectives, and digital images obtained with DP72 CCD or DP73 cameras (Olympus) using CellSens Standard software (Olympus).

For establishing the platform, we used a custom designed PCD siRNA library (Dharmacon) comprising 81 siRNAs targeting apoptotic, necroptotic and autophagic genes. These were arranged randomly in two 96 well white flat bottom plates (Greiner, 655083) with several control siRNAs scattered among them (25 nmol each). Eight siRNAs were placed on both plates as internal standards to enable plate comparison. The melanoma cell lines were then reverse transfected in the 96 well plates (5000 cells/well). 24h post-transfection the medium was replaced with fresh medium, and 48h post transfection vemurafenib (10μM) was added to the treated cells and DMSO to the untreated cells. After 24h of treatment, cell viability was measured by the CellTiter-Glo assay. Each cell line was assessed in triplicate, on different days.

Analysis of the functional death signature was done using R as follows: The value of the 8 genes that were represented on both plates was the average of the two technical repetitions. From the raw values, the value of blank cells was subtracted, then the average value of the si-controls from each plate was used to normalize the value of each siRNA. The si-control value was set as 1. The normalized values from each repetition were averaged. The ratio of the ln average of the treated divided by the ln average of the untreated value was the mean ratio that was used to determine the change in the viability. A value of 0 refers to no change in the viability when knocking down the gene between the treated and the untreated samples, positive and negative values refer to increased and decreased viability, respectively. Hits were those knock-downs that passed two criteria: the change in the viability had to be significant (*p*<0.05) in comparison to the si-control according to a Dunnet test, and in comparison to the untreated KD with the same siRNA by student T-test. Both *p*-values appear in Supplementary Table S2.

### Cell cycle analysis

For cell cycle analysis, 2×10^6^ cells were pulsed with 10μM BrdU (Sigma, B5002) for 2-4h and both detached cells and trypsinized attached cells were collected. Cells were fixed in cold ethanol overnight at 4°C, incubated for 30 min in Denaturation Solution (2N HCl/Triton X-100 in PBS 1X), washed in Neutralization Solution (0.1 M Na_2_B_4_O_7_, pH 8.5) and incubated 1h with Antibody Solution (PBS 1X, 0.5% Tween 20, 1% BSA, 1μg/100ul anti-BrdU FITC antibody (eBioscience, cat#11-5071-41). Cells were then resuspended in PBS containing 50 μg/ml PI and 50 μg/ml RNase A. Flow cytometry analysis was performed on a BD LSRII instrument (BD Immunocytometry Systems) equipped with 355-, 407-, 488-, 561- and 633-nm lasers, controlled by BD FACS Diva software v8.0.1 (BD Biosciences), at the Weizmann Institute of Science's Flow Cytometry Core Facility. FITC was detected by excitation at 488 nm and collection of emission using 505 longpass (LP) and 525/50 bandpass (BP) filters. PI was detected by excitation at 561 nm and collection of emission using 600 LP and 610/20 BP filters. Further analysis was performed using FlowJo software v10.2 (Tree Star).

### Protein analysis

Cells were lysed in PLB buffer (10mM NaPO_4_ pH 7.5, 5mM EDTA, 100mM NaCl, 1% Triton X-100, 1% Na deoxycholate, 0.1% SDS) supplemented with 10μl/ml 0.1M PMSF and 1% protease and phosphatase inhibitor cocktails. Proteins were separated by SDS-PAGE and transferred to nitrocellulose membranes, which were incubated with antibodies against ERK (cat#M5670), pERK (cat#M8159), vinculin (cat#V9131) (Sigma) FIP200 (cat#12436), Atg12 (cat#4180P), BIM (cat#2819), cleaved caspase-3 (cat#C-9664) (Cell Signaling), or Mcl-1 (Santa Cruz, sc-20679). Detection was done with either HRP-conjugated goat anti-mouse or anti-rabbit secondary antibodies (Jackson ImmunoResearch cat#211-032-171, 111-035-144), followed by enhanced chemiluminescence (EZ-ECL, Biological Industries Israel Beit-Haemek Ltd.)

### Statistics

The statistical significance of differences between means was assessed either by standard Student’s T-test or by one-way ANOVA followed by Tukey’s post hoc test, as stated in the figure legends. Values with *p*<0.05 were considered significant. Statistical analysis was done using GraphPad Prism version 7.04 (La Jolla California, USA) or for the large screen viability output, R 3.3.1 (2016-06-21), with the nparcomp, multcomp and the pheatmap packages.

## Results

### Patient-derived melanoma cell lines carrying the BRAF V600E mutation show heterogeneous responses to vemurafenib treatment

A cohort of 12 early passage melanoma cell cultures were established from individual patient tumors carrying the V600E mutation of *BRAF*. Tumour cell lines were derived from metastatic patients before treatment with the clinically approved BRAF inhibitors was available (Supplementary Table S1). Despite the common activating *BRAF* mutation, there was a wide heterogeneity in the response of the cell lines to 24h treatment with 10μM vemurafenib; the extent of reduction in cell number/viability, as assessed by the CellTiter-Glo assay, ranged from 20-80% in 11/12 cell lines, and one cell line showed no change (Fig. 1A). Western blotting indicated a decline in phosphorylation of ERK, the downstream target of BRAF, in all cell lines except for line 13, indicating that the drug was effective in inhibiting BRAF in the other 11 cell lines (Fig. 1B). Notably, in line 13, the high basal ERK phosphorylation levels were even further increased by the drug, suggesting that ERK phosphorylation is upregulated in this tumor by some second unknown mechanism independent of the V600 *BRAF* mutation. BIM (BCL2L11) protein levels were elevated in all drug-sensitive cell lines, except for line 43, a consequence of its stabilization upon reduced phosphorylation by ERK (23).

**Figure 1.**
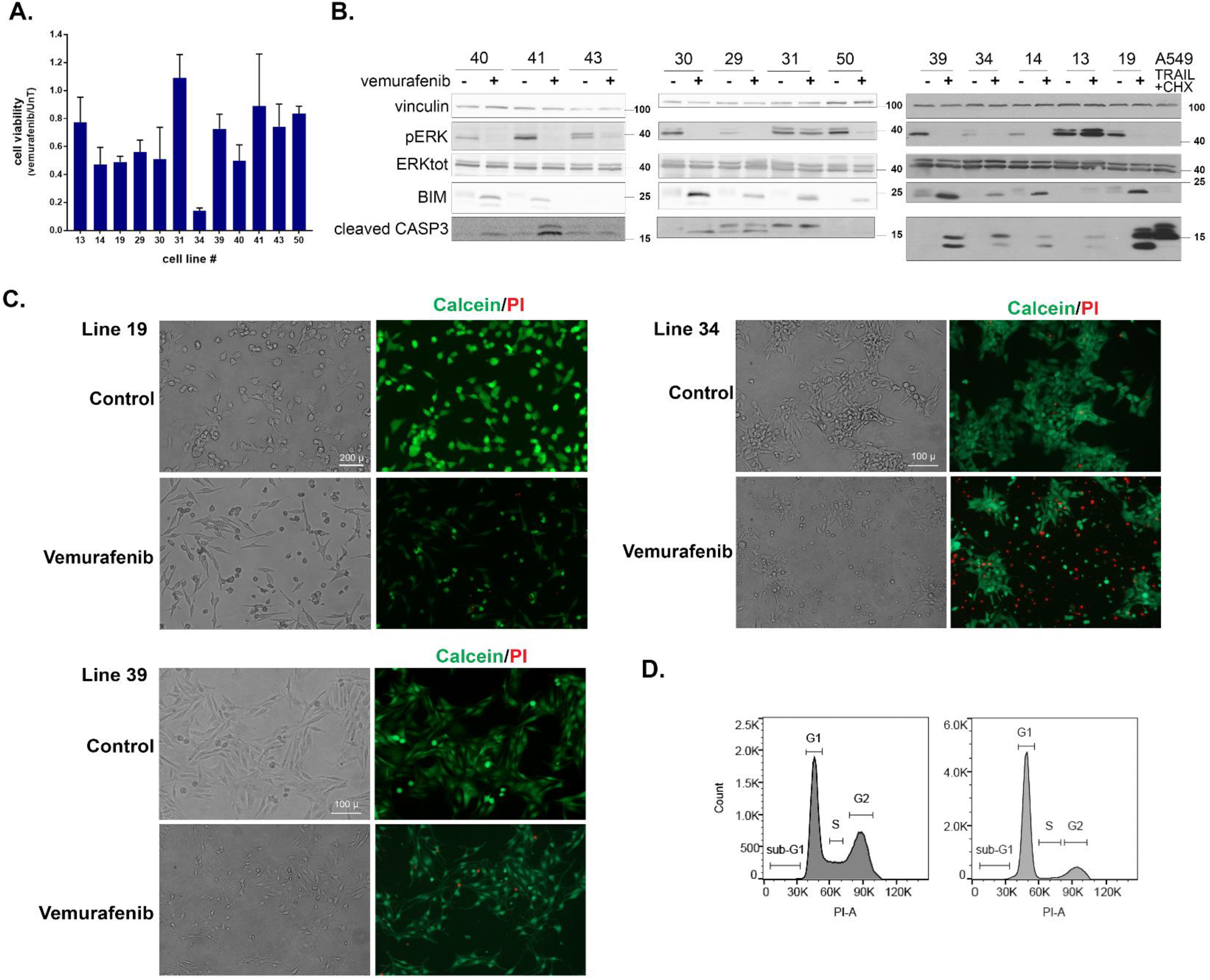
Patient-derived melanoma cell lines show heterogeneous responses to vemurafenib treatment. **A.** Quantification of cell viability by CellTiter-Glo assay in response to 10μM vemurafenib treatment for 24h. CellTiter-Glo reads of the untreated cell lines (UnT) were normalized to a value of 1 and the y-axis values indicate the relative decrease in response to vemurafenib treatment. Shown are the mean±SD of triplicates, for each cell line. **B.** Western blots monitoring the responses of the 12 melanoma cell lines to vemurafenib treatment showing: ERK phosphorylation, Bim expression levels and cleaved Caspase-3 levels. A549 cells treated with TRAIL and cycloheximide were used as a positive control for strong caspase-3 activation during apoptosis. Lysates were run on three different blots as the cell lines became available for analysis, and blots were cut into appropriate strips for simultaneous probing with antibodies to proteins of different sizes. Antibodies to vinculin were used as loading control on all blots. For ERK detection, blots were first probed for phospho-ERK, stripped and reprobed with antibodies to total ERK. For the blots of cell lines shown at left and right, the same lysate was run on two separate gels to examine proteins with overlapping sizes, for the middle set of cell lines, only 1 gel was run, and the slice of membrane used to detect Caspase-3 was stripped and reprobed with anti-Bim antibodies. (Uncropped version of these blots can be seen at end of Supplementary Data*, for reviewers only*). **C.** Representative phenotypic response to vemurafenib of cell lines 19, 34 and 39. Cells were treated with vemurafenib for 24h, and then stained with calcein AM (green, stains live cells) and propidium iodide (PI) (red, stains cells with permeabilized membranes) and viewed at 10x (lines 34, 39) or 20x magnification (line 19). Greyscale versions of single filter channels for calcein and PI can be seen in Supp. Fig. S1A. **D.** Cell cycle distribution of line 39 treated with 5μM vemurafenib for 24h and stained with PI. 50,000 cells were recorded for each sample. Shown is a representative experiment of three independent repetitions. The X-axis corresponds to PI staining and the Y-axis to cell number.

Phenotypic analysis following staining with calcein-AM/PI indicated that the reductions in CellTiter-Glo values were caused by at least three different cell responses during the first 24h of drug treatment (Fig. 1C and Supplementary Fig. S1A). A high incidence of dying round, shrunken cells was observed in some cell lines, such as lines 34 and 19. This was associated with membrane permeabilization, as documented by the strong PI staining, in line 34, but not in line 19, suggesting that line 34 cells die by programmed necrosis, whereas line 19 cells undergo an early apoptotic response. In contrast, in cell line 39, the abundance of cells with a death related phenotype at the 24h time-point was low, suggesting that the reductions in the CellTiter-Glo values mostly reflect proliferation arrest rather than cell death. Flow cytometry cell cycle analysis of line 39 confirmed the microscopic observations. At 24h post drug treatment, the cells accumulated in G1, at the expense of S and G2 phases (Fig. 1D), and had reduced BrdU incorporation (Supplementary Fig. S1B), indicative of a block in cell cycle progression. This initial morphological survey documents heterogeneity in drug responses to vemurafenib among patient-derived cell lines ranging from cytostatic and different types of cytotoxic responses.

Molecular analysis of the death response by western blotting was consistent with the observed phenotypic heterogeneity (Fig. 1B). Caspase-3 (CASP3) was cleaved to its activated products to varying extents in approximately half of the cell lines, with line 19 showing the most robust cleavage, similar to the positive control of A549 cells treated with TRAIL and cycloheximide (Fig. 1B). The remaining cell lines either failed to activate caspase-3, or did not show enhanced caspase-3 cleavage in response to the drug compared to control. This was despite the suppression of the MAPK pathway and stabilization of the pro-apoptotic BH3-only protein BIM in all but one of these cell lines. This suggests that BIM stabilization is not sufficient in itself to promote caspase-activating pathways. Thus in these latter cell lines, caspase-independent pathways are limiting the growth and/or cell viability, at least during the first 24h of drug treatment. Altogether these initial experiments underscore the large heterogeneity that exists in the phenotypic and molecular responses to vemurafenib treatment in melanoma.

### Mapping points of vulnerability in the programmed cell death map of patient-derived melanoma cell lines

The less than optimal and variable death responses to vemurafenib within the melanoma cohort suggest that strategies to further reduce cell viability should be “personalized” to activate the most effective cell death pathway in each cell line. To this end, based on the PCD map that was curated in our lab (Supplementary Fig. S2A), we generated a siRNA-based pre-screening platform for studying the fidelity of various cell death pathways in individual melanomas, to identify soft-spots in the cell death machinery that can be targeted for maximal killing in each tumor. The detailed outline of the platform is depicted in Fig. 2A. Briefly, a custom designed PCD siRNA library (Dharmacon) comprising 81 siRNAs targeting apoptosis, autophagy and programmed necrosis genes were arranged randomly in two 96-well plates with several control siRNAs scattered among them (Supplementary Fig. S2B). The siRNAs were reverse transfected into the melanoma cell lines, and cell viability in response to vemurafenib was then assessed for individual knock-down cells using the CellTiter-Glo assay. For each cell line, the screen was done in triplicate (see Supplementary Table S2 for the output). These results were converted into heat maps depicting the varying levels of viability for each KD cell line, and were organized in a combined global heat map display for the 12 cell lines (Fig. 2B). A functional hit was defined as a statistically significant siRNA-mediated effect that moderated the vemurafenib-induced reduction in cell viability, by either enhancing cell death (positive hits, red bars of varying intensity in Fig. 2B) or by decreasing cell death (negative hits, blue bars of varying intensity in Fig. 2B). By determining the contribution of each gene to the death response to vemurafenib in this manner, the overall ‘functional death signature’ of each tumor was characterized, highlighting the specific death pathways that are functionally active. The positive hits identified within these pathways are the ‘soft-spots’ that can be targeted for enhanced cell killing.

**Figure 2.**
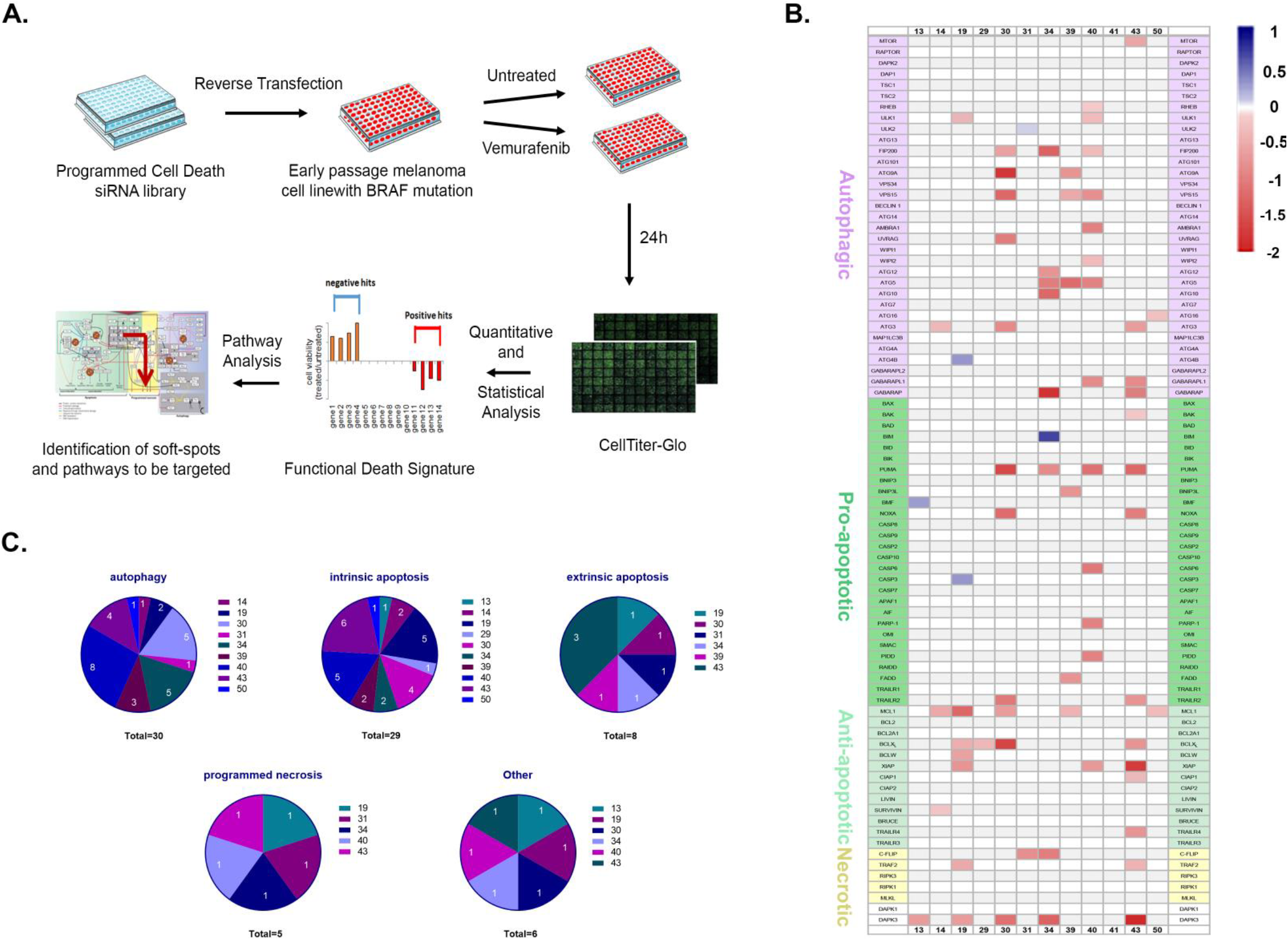
Mapping points of vulnerability in the programmed cell death map in each tumor. **A.** Schematic of the siRNA-based screening platform used for delineating the functional death signature of each cell line. Each experiment was done in triplicate. **B.** Combined heatmap of the hits for all cell lines. Red corresponds to positive hits (soft-spots), blue to negative hits, and genes whose knock-down did not statistically affect the responses to the drug are in background color (white or light grey). Genes are grouped according to cell death sub-modules; DAPK1 and DAPK3, multifunctional regulators of various pathways that are not part of the basic cell death machinery, are not included in any specific sub-module. **C.** Pie charts indicating distribution and number of soft-spots per cell line, within each cell death sub-module (autophagy, intrinsic and extrinsic apoptosis and programmed necrosis; “other” refers to the DAPK family, specifically DAPK3, the only soft-spot within this group).

The global overview of the combined heat maps of the 12 cell lines showed a large heterogeneity in the number and position of the soft-spots within the three modules of the PCD map (Fig. 2B and Supplementary Table S3). The average number of soft-spots was 6, however in some cell lines (19, 30, 34, 39, 40, 43) there were up to 15 soft-spots scattered among the PCD modules, mostly targeting different genes along the PCD pathways in each cell line, with little overlap (see Supplementary Fig. S3 for positioning of the hits in the PCD map in lines 19, 34 and 39). The other cell lines displayed fewer soft-spots. Cell lines 13, 29 and 31 each displayed a single soft-spot at different positions in the PCD map, and cell line 41 had no detectable soft-spots.

Several interesting conclusions emerged from this analysis. Not surprisingly, the intrinsic apoptosis pathway was important for enhancing the cell death response to vemurafenib (Fig. 2C); soft-spots corresponding to apoptosis inhibitors from within this pathway were observed in 10/12 cell lines. Importantly, the identities of the soft-spots differed among the cell lines, and mapped to various members of the BCL2 and/or IAP families. The most common positive hit was the anti-apoptotic BCL2 family member MCL1, which was found in 5/12 cell lines. Counterintuitively, in 4 cell lines, KD of PUMA (BBC3) or NOXA (PMAIP1), members of the BH3-only protein subfamily, led to enhanced killing. This suggests that in specific cellular contexts, these proteins may display a function opposite of the canonical pro-apoptotic roles previously reported for BH3-only proteins. Hits from the extrinsic apoptotic pathway were also identified in 5 cell lines (Fig. 2C); of these, only line 31 had no hits within the intrinsic pathway.

The second module that emerged with multiple soft-spots was autophagy (Fig. 2C). Clusters of positive hits within the various steps of autophagy were found in 5/12 cell lines, suggesting that autophagy serves as a survival mechanism in these drug-treated melanoma. For example, cell line 34 had 5 autophagy soft-spots (ATG10, ATG12, ATG5, FIP200 (RB1CC1) and GABARAP) (Fig. 2B and Supplementary Table S3, Supplementary Fig. S3B). Four of these five genes belong to the ubiquitin-like conjugation pathway that mediates LC3B (MAP1LC3B) conjugation to phosphatidylethanolamine (PE), representing the central execution step of autophagy, membrane elongation. In other cells (lines 30, 39, 40, 43), the hits were more widely dispersed among the different steps. ATG5 and FIP200 were the most common hits, found in 3 cell lines each. A second profile of autophagy soft-spots was also observed (Fig. 2C): three cell lines (lines 14, 19, 50) had single hits at different positions within the pathway, as opposed to multiple hits throughout the pathway (see for example line 19, Supplementary Fig. S3A). In these cell lines, the emergence of lone autophagy proteins as soft-spots suggests that these genes may serve a second cell death function independent of autophagy, a phenomenon that has been documented for several autophagy proteins (21) and is depicted by cross-pathway interactions within our PCD map.

Interestingly, the most frequent soft-spot was the Ser/Thr kinase DAPK3 (positive hit in 6/12 cell lines) (Fig. 2B,C and Supplementary Table S3).). Knock-down of DAPK3 was validated in cell line 34 (Supplementary Fig. S4A). In 4 additional cell lines, KD of DAPK3 alone significantly reduced cell viability and the addition of vemurafenib did not enhance cell killing, and therefore it did not pass our definition of a positive hit (Supplementary Table S2).

In order to validate the large scale screen results, and study the effect of the knock-down on other critical molecular responses to the drug, two cell lines with multiple hits were further analysed. In cell line 34, knock-down of ATG5 reduced its mRNA expression and the steady state levels of the ATG5-ATG12 conjugate (Supplementary Fig. S4B,C), confirming that the ATG5 siRNA is effective in limiting the conjugation process with ATG12, a step that is critical for autophagosome maturation. FIP200 protein levels were also reduced by the corresponding siRNA to FIP200 (Supplementary Fig. S4C). Notably, the knock-down of these two autophagy genes did not affect the upstream dephosphorylation of ERK and consequent stabilization of BIM in response to vemurafenib treatment, suggesting a downstream effect. Also their knock-down did not accelerate the low levels of caspase-3 activation by vemurafenib characteristic of this cell line, implying that the increased cell lethality that resulted from autophagy perturbation did not involve the indirect activation of the apoptotic module. As shown in Fig. 3A, the KD of ATG5 and FIP200, and also ATG12, DAPK3 and C-FLIP (CFLAR) (the other three soft-spots identified in this cell line) reduced the overall cell viability in response to vemurafenib, similar to the original screen, thus validating the screen results for this cell line.

**Figure 3.**
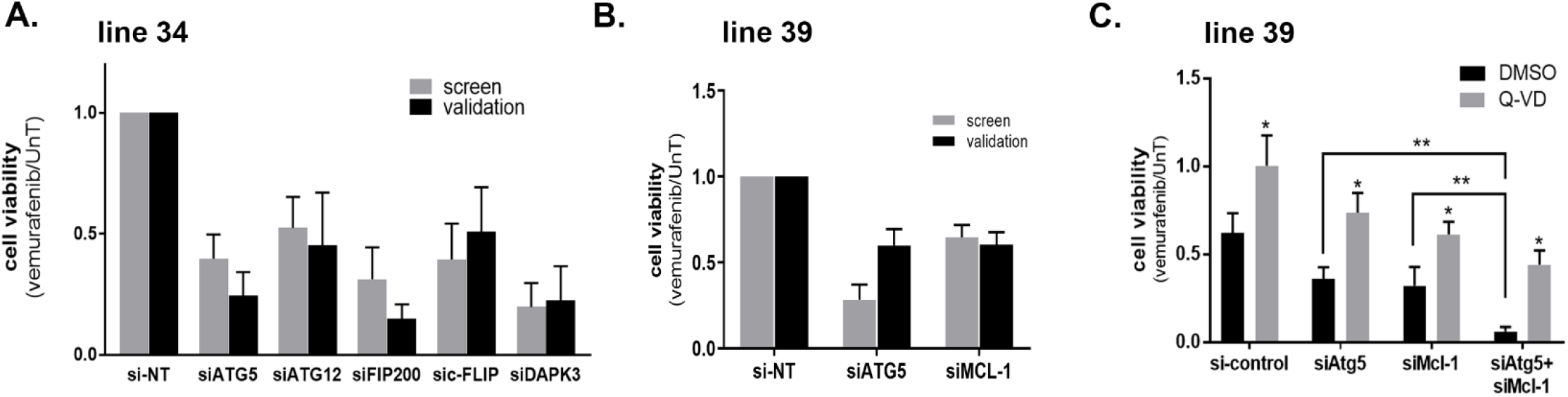
Validating points of vulnerability in the programmed cell death map in each tumor. **A.-C.** Cell viability in response to 10μM vemurafenib was tested 48h after transfection with the indicated siRNAs or siRNA combinations using the CellTiter-Glo assay in line 34 (A) or line 39 (B,C). In C, 50μM Q-VD, or DMSO as control, were added. The values were first normalized to si-control knock-down cells, and then the ratio of drug treated to untreated calculated, with the ratio of the si-control knock-down cells set as 1. Data represents mean±SD of three replicate experiments. In C, statistical significance of the difference across siRNA manipulations was assessed using one-way analysis of variance (ANOVA) followed by Dunnett's multiple comparisons test, ** *p* <0.01. Statistical significance of Q-VD vs. DMSO treatment was determined by two-tailed paired Student’s T-test, * *p*<0.05.

Cell line 39 had 6 hits (Fig. 2B and Supplementary Table S3, Supplementary Fig. S3C); validation assays were performed for MCL1 and ATG5 siRNAs. The cell death responses of the KD in combination with vemurafenib were similar to those observed in the large-scale siRNA screen (Fig. 3B). As in cell line 34, the KDs did not interfere with upstream dephosphorylation of ERK and stabilization of BIM as a response to vemurafenib treatment (Supplementary Fig. S4D). Notably, the KD of anti-apoptotic MCL1 increased cleaved caspase-3. Similarly, and in contrast to line 34, ATG5 KD also led to enhanced caspase activation. The effects of both KDs were partially rescued by the caspase inhibitor Q-VD (Fig. 3C). This suggests that the knock-down of ATG5 in combination with vemurafenib augments activation of the apoptosis pathway, supporting functional duality. Interestingly, the combined knock-down of ATG5 and MCL1 had an additive effect (Fig. 3C), suggesting that they most likely function via different mechanisms to inhibit apoptosis, and that apoptosis can be enhanced by more than one pathway. The additive effects of double perturbations underscore the importance of unveiling several potential non-redundant soft-spots in each tumor.

### Inhibition of autophagy by drugs promotes cell death induced by vemurafenib in a cell line specific manner

In order to further translate the identified tumor points of vulnerability towards possible clinical directions, siRNAs corresponding to positive hits were replaced with clinically approved drugs, when available. In cell line 34, the identification of several autophagy genes as soft-spots that are clustered along the ubiquitin-like conjugation pathways (ATG5, ATG12, ATG10, GABARAP) suggested that autophagy as a whole process was critical for counteracting the cell death responses to vemurafenib. Thus, available autophagy inhibitors were used and their effects were compared to ATG5 knock-down. Treatment with the combination of vemurafenib and lysosomal inhibitors hydroxychloroquine (HCQ) or Bafilomycin A, which block fusion of the lysosome with the autophagosome, enhanced the reduction in cell viability compared to vemurafenib alone by 50% (Fig. 4A). Neither autophagy inhibitor had any effect on cell viability alone (Supplementary Fig. S4E). In contrast, cell line 39, which had only 3 sporadic autophagic hits, at least one of which (ATG5) was linked to the apoptosis pathway, did not show enhanced cell killing when treated with HCQ or Bafilomycin A in combination with vemurafenib (Fig. 4B). Thus the autophagy pathway is a promising target for increasing vemurafenib-induced death responses only if the siRNA-based pre-screening platform indicates multiple potential soft-spots that cluster to the main autophagy pathway stages.

**Figure 4.**
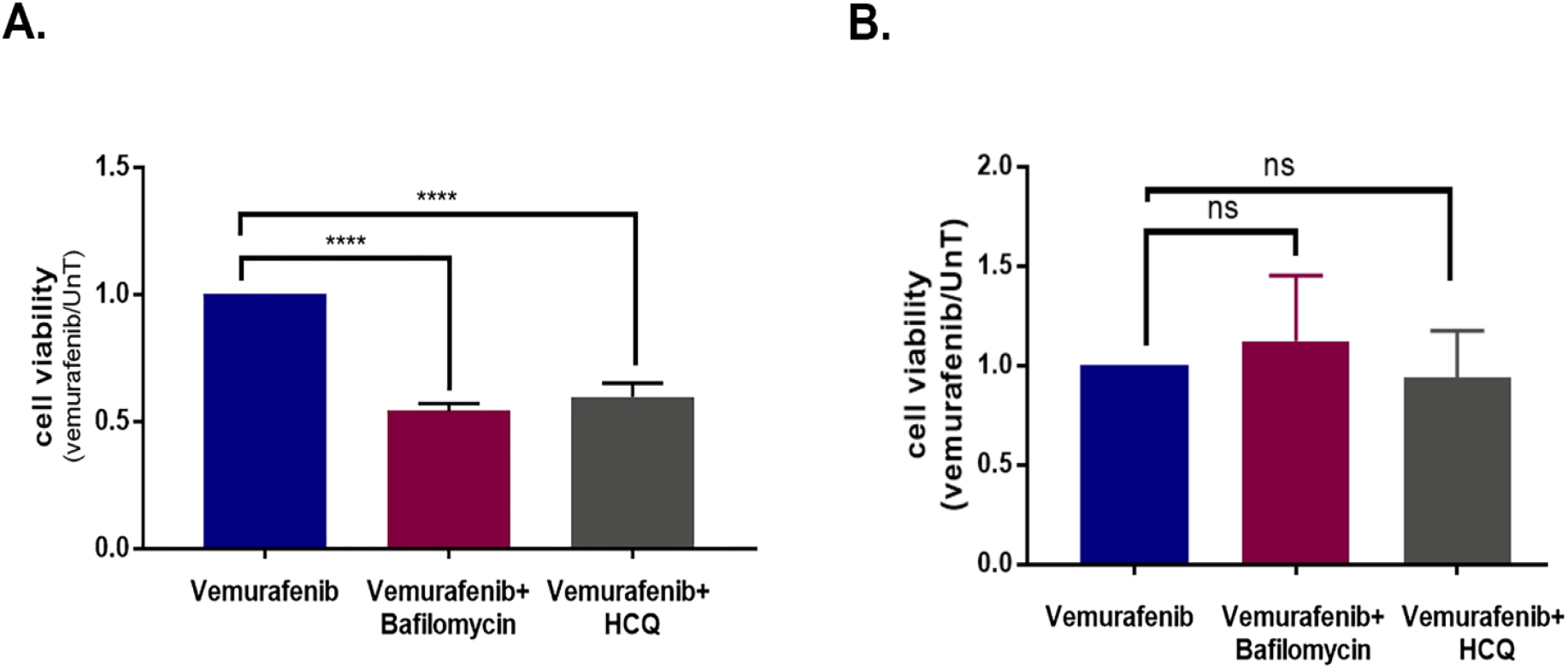
Inhibition of autophagy promotes cell death induced by vemurafenib in a cell-line specific manner. **A., B.** Cell viability of cell lines 34 (A) and 39 (B), treated with the combination of 10μM vemurafenib and the autophagy inhibitors Bafilomycin A1 (5nM) or HCQ (20μM) for 48h, as assessed by CellTiter-Glo assay. Data represents mean±SD of three replicate experiments. Statistical significance was assessed by ANOVA followed by Dunnett's multiple comparisons test, **** *p*<0.0001.

### Inhibition of BCL2 family members by clinically approved drugs promotes cell death induced by vemurafenib in a cell line-specific manner

Not surprisingly, the anti-apoptotic BCL2 family members emerged as soft-spots in multiple cell lines, with MCL1 the most prominent among them, although different cell lines had different signatures for this protein family (Fig. 5A). For further investigation, the various knock-downs were replaced with specific inhibitors S63845 for MCL1, A1331852 for BCLX_L_ (BCL2L1), venetoclax for BCL2, and the more wide-range ABT-737, which targets BCL2, BCLX_L_ and BCLW (BCL2L2), in cell lines 14, 19, 39, 40, and 43 (Fig. 5A). For each cell line, there was a significant reduction in the viability of the cells after 48h co-treatment with vemurafenib, only upon inhibition of the BCL2 family members that were positive hits in the screen. For example, in line 39, the MCL1 specific inhibitor S63845, but not the BCL2 or BCLX_L_ specific inhibitors, led to an enhanced cell death response to vemurafenib (Fig. 5B). S63845 also enhanced cell killing in lines 14 and 19, in which MCL1 was a soft-spot, as in line 39 (Fig. 5C, D). In contrast, the MCL1 inhibitor had no additional effect on cell killing in line 43 (Fig. 5E), in which MCL1 was not a positive hit, although the BCLX_L_ inhibitor did enhance cell killing. The broad range ABT-737 that does not inhibit MCL1 had no effect on line 39 or line 14, but did reduce cell viability in line 19, in which BCLX_L_ and BCLW were identified as positive hits (Fig. 5B–D). Notably, none of the BCL2 family inhibitors had any effect on vemurafenib-induced cell killing in line 40, which showed no soft-spot within the family, and served as the negative control in these experiments (Fig. 5F). There was no change in viability from treatment with inhibitors alone (see for example, Supplementary Fig. S4E).

**Figure 5.**
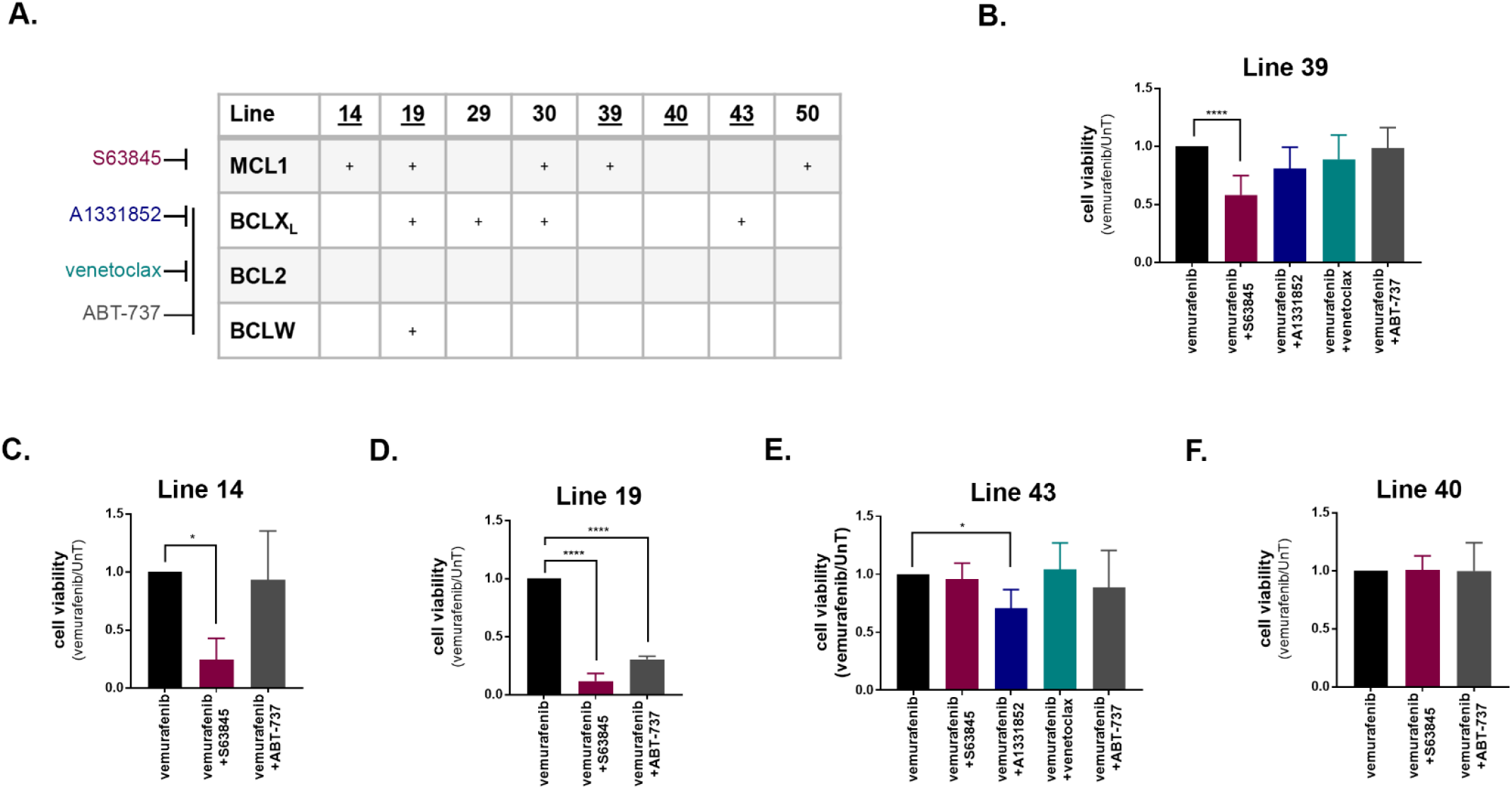
Inhibition of BCL2 family members promotes cell death induced by vemurafenib in a cell-line specific manner. **A.** Chart indicating positive hits within the anti-apoptotic BCL2 family in the indicated cell lines. Specificity of inhibitors are indicated at left. **B.-F.** Cell viability of cell lines treated with the combination of 10μM vemurafenib and either 0.5μM (B-D,F) or 1μM (E) of inhibitors of BCL2 family members for 48h. Data represents mean±SD of three replicate experiments. Statistical significance was assessed using ANOVA followed by Dunnett's multiple comparisons test, * *p*<0.05, **** *p*<0.0001.

### Targeting soft-spots during long term vemurafenib treatment of melanoma cell lines reduces viability of drug tolerant cells and the number of drug resistant foci

The most important clinically relevant question is whether targeting the soft-spots identified in the screen affects long term vemurafenib treatment, and whether it translates into reduced frequency of acquired drug resistance. To assess this, the additive effects of S63845 on the vemurafenib responses in cell line 39 were measured by flow cytometry for DNA content and by CellTiter-Glo for cell viability over a 14d time-course. While treatment with 2μM vemurafenib alone induced G1-arrest during the first 24h (Supplementary Fig. S5A), the combination of 2μM vemurafenib and 0.5μM S63845 showed an apoptotic response, as indicated by the elevated sub-G1 population (Fig. 6A, Supplementary Fig. S5A). At later time-points, the sub-G1 population was always larger in the cells that were treated with the drug combination (Fig. 6A), indicative of accelerated apoptosis. The cell viability assay gave similar results; from 1d and up to 14d of treatment, the viability of the cells treated with the drug combination was significantly lower than the viability of cells treated with vemurafenib alone (Fig. 6B). Notably, S63845 alone had no detectable effects, as measured both in the cell viability assay (Fig. 6B) and by flow cytometry for DNA content (Supplementary Fig. S5A).

**Figure 6.**
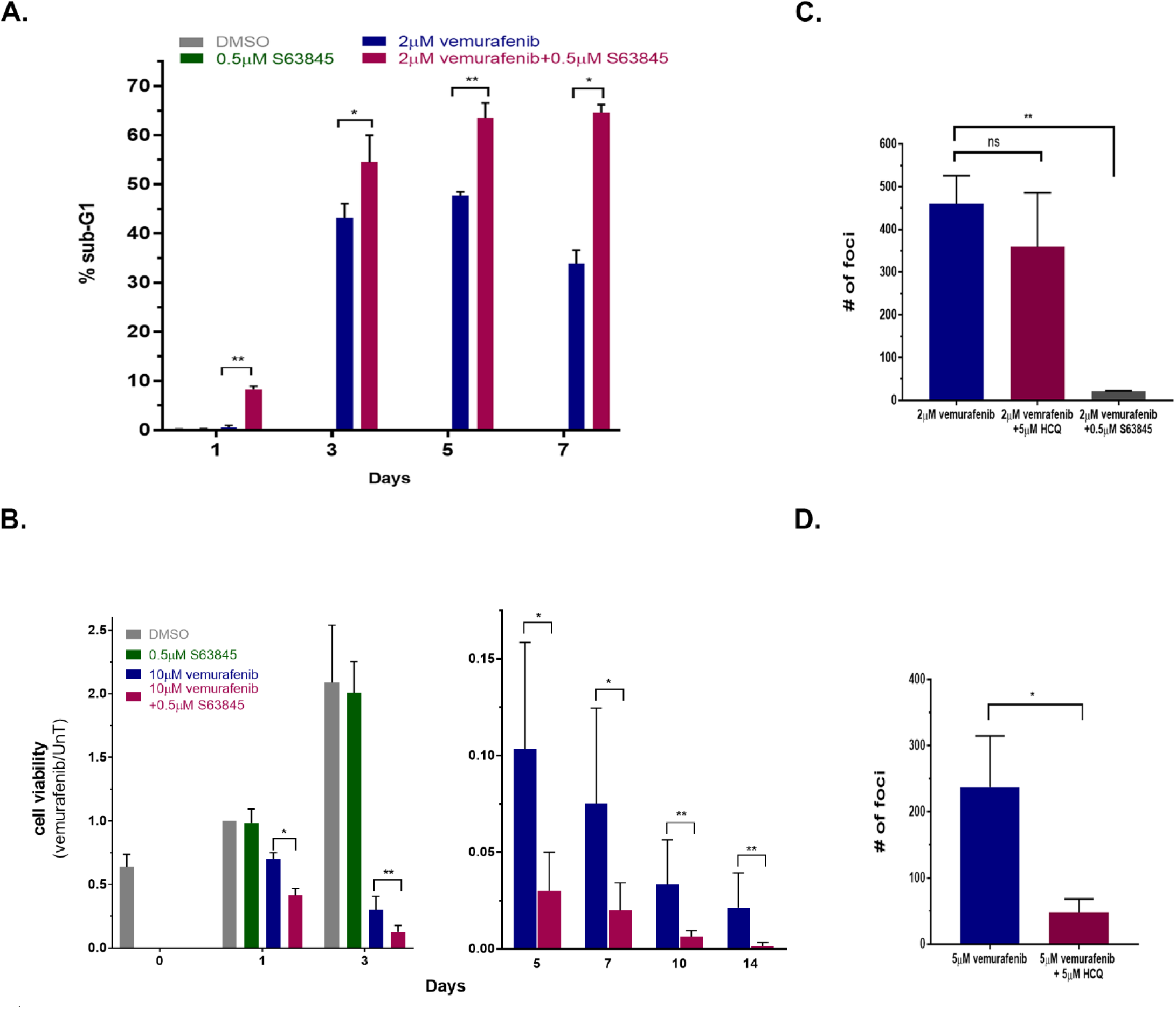
Targeting MCL1 during long term vemurafenib treatment of melanoma cell lines reduces viability of drug tolerant cells and the number of drug resistant foci. **A.** Line 39 cells were either untreated, or treated with 2μM vemurafenib, 0.5μM S63845 or both for 1, 3, 5 and 7 days. Cells were stained with PI and 50,000 cells per sample were analyzed by FACS in 3 replicates. Shown is the % sub-G1 population. Cells that were treated with DMSO or S63845 alone were only assessed at day 1 since they reached confluence beyond that point (0.18% and 0.28%, respectively). * *p*<0.05, \*\**p*<0.01. **B.** Cell viability of line 39 left untreated or treated with 10μM vemurafenib, 0.5μM S63845 or both for up to 2 weeks, as assessed by CellTiter-Glo assay. Monitoring of untreated and S63845 treated control cells was discontinued after 3 days, and the data is split accordingly between two graphs with different Y-axis ranges. Values were normalized to day 1 untreated. **C, D.** 6×106 cells of line 39 (C) or line 34 (D) were plated and treated with 2μM vemurafenib alone or with 5μM HCQ or 0.5μM S63845 for 4 and 5 weeks, respectively. Fresh medium containing the drugs was supplied twice weekly. Individual foci were counted, shown are mean±SD of three independent biological repeats. The treatment with either 0.5μM S63845 or 5μM HCQ alone did not have an effect on the cells, comparable to no treatment, which reached full confluence by 3 days, and thus foci could not be counted within the same time-course and plating conditions. \*\**p*<0.01, ns- non-significant.

Under the continuous vemurafenib treatment, a small fraction of drug tolerant cells was identified displaying the main characteristics of persister cells. That is, the population of surviving cells retrieved and grown after removing the drug at day 10 of treatment retained their sensitivity to vemurafenib similar to the original naïve population (Supplementary Fig. S5B). The number of drug tolerant cells that still maintained their viability was further reduced as drug treatment continued (Fig. 6B), yet this fraction of cells was not completely eliminated. In contrast, the combination of vemurafenib and MCL1 inhibitor was consistently more effective in reducing cell number during prolonged treatment. After several weeks of continuous exposure to vemurafenib, drug resistant colonies (foci) developed in the plates from the residual cell population. Resistant colonies started to appear in lines 39 and 34 after 35 days of treatment. A cluster of cells was considered to be a focus only when mitotic cells were abundantly identified within it. Most importantly, in line 39, the number of drug resistant foci was reduced 22-fold in the plates simultaneously treated with the MCL1 inhibitor S63845 and vemurafenib, as compared to vemurafenib alone (Fig. 6C). In contrast, the combination of vemurafenib and HCQ did not change the number of drug resistant foci, consistent with the inability of the autophagy inhibitor to enhance the initial responses to vemurafenib (Fig. 3B) and with the functional death signature of line 39 in the screen (Fig. 2B; Supplementary Fig. S3C). When testing the combination of vemurafenib with or without HCQ in cell line 34, where the autophagy pathway was identified as a point of vulnerability, the combination treatment reduced foci formation five-fold (Fig. 6D). In conclusion, the combination of vemurafenib and drugs that targeted the soft-spots identified by the siRNA screen reduced the number of viable cells under continuous drug treatment, and as a consequence, reduced the number of resistant colonies emerging from this residual cell population.

## Discussion

Genomic and transcriptomic tools can predict which oncogenes drive a particular cancer, enabling the design of targeted treatment. However they cannot capture the full genetic and epigenetic alterations and complexity of the tumor, and patients that are given targeted therapy solely based on their -omics data tend to relapse (24,25). Drug combinations are needed to reduce the number of cells that survive below the critical number that enables the development of acquired drug resistance with time. Therefore, functional screening approaches were developed using siRNA and drug libraries in cell lines and organoids obtained from patients to identify novel combinations (26–30). Montero, et al. (30) developed a functional screen, called Dynamic BH3 Profiling, that can predict if a tumor is primed for apoptosis after certain chemotherapy, based on their response to a panel of BH3 mimetics. In the present study, we undertook a more global approach to profile the broader functional cell death network for personalized precision therapy for melanoma. An siRNA screening platform was used to determine the functional death signature of naïve melanoma tumors carrying V600E *BRAF* mutation challenged in culture with vemurafenib. This enabled the identification of points of vulnerability within the cell death network, referred herein as soft-spots, which can lead to therapeutic strategies using drug combinations tailored to each patient to initiate a strong death response. We have shown that such combinatorial approaches enhance cell death and most importantly, reduce the emergence of resistant clones during long-term growth in culture. Notably, the soft-spots that emerged from the siRNA screen comprised only those targets that were not lethal in the absence of vemurafenib, thus conferring selectivity of the combinatorial treatment towards the tumor cells carrying the *BRAF* mutation. The ability of this pre-screening platform to map various hits along the apoptosis, autophagy and programmed necrosis pathways provides a broad view of the functional death signature of each tumor, and opens the opportunity for simultaneously targeting multiple hits to further increase the lethal responses, as shown here by the double perturbations of ATG5 and MCL1 in line 39. Clinically, this approach would translate to a personalized treatment protocol aimed at decreasing the tumor size below the threshold that allows the tumor to acquire drug resistance and prevent relapse of the tumor.

Our analysis demonstrates the necessity of mapping the intact cell death pathways that can be activated in each individual tumor. The molecular and phenotypic analysis of the twelve early passage melanoma cell lines derived from individual patient tumors showed different types of responses to vemurafenib treatment, and the siRNA perturbations indicated different functional death signatures in each cell line. This could reflect the fact that the BRAF-ERK1/2 signaling axis can be connected to different cellular processes, and thus its inhibition will have divergent effects on pathways controlling cell growth and death, both caspase-dependent and independent.

It was not surprising that many apoptotic regulators emerged as soft-spots, as vemurafenib treatment has been documented to activate apoptosis in many cases (31,32). Yet both the diversity of the soft-spots, and the unexpected emergence of some pro-apoptotic proteins among them, underscore the importance of conducting unbiased personalized screens to identify the best means of activating the apoptosis module during vemurafenib treatment. MCL1 and BCLX_L_ were the BCL2 family members present as the most frequent soft-spots, consistent with other studies that have documented high levels of MCL1 and BCLX_L_ in melanoma (33–36) and that have shown that combination treatment of BRAF inhibitor with BH3-mimetics was effective in melanoma cell lines (37,38). However, choosing the right inhibitor that would be most effective is still a challenge; the analysis presented here shows that different melanomas respond differently to the range of BH3 mimetics currently available. The siRNA pre-screening platform can help to determine when and which BH3-mimetic should be administered.

In addition to apoptosis, autophagy was also found to play a role in the survival of melanoma cell lines. Successful combination of inhibitors from the MAPK pathway with autophagy blockers has been documented in several cancers, such as brain tumors (39), pancreatic cancer (40) and melanoma (16,41,42). This led to the suggestion that autophagy helps cells cope with the stress induced by drug treatment, and therefore blocking autophagy removes this coping mechanism and enhances the death response. Most of the autophagic hits fell within the ubiquitin-like conjugation pathway that executes the membrane elongation steps of autophagy. Upstream regulators were less prominent; this may be due to the abundance of signaling pathways that converge on the initiation stages of autophagy, redundancy among these regulators (e.g. ULK1 and ULK2), the multiple sources for phagophore membrane initiation, and/or the presence of non-canonical signaling pathways that can occur independently of the VPS34 (PI3KC3) PI3K lipid kinase and ULK1 complexes. The ubiquitin-like conjugation step, in contrast, may serve as a limiting process for the continuation of autophagy. Of note, autophagy inhibitors in use in the clinic to date, such as chloroquine (CQ) and HCQ, are not specific for the autophagy process. Pre-clinical attempts to develop more specific inhibitors have focused on kinases such as VPS34 (43). Our study, however, indicates that these upstream targets may not be rate-limiting and thus not the best candidates for blocking this survival pathway in melanoma, but rather, drug design should focus on the ubiquitin-like conjugation step. As in the case of the BCL2 family inhibitors, the cell death signature data indicates that inhibiting autophagy is not effective in all melanoma tumors; our study suggests that pre-screening tumor responses to vemurafenib can predict the effectiveness of autophagic inhibitors as a viable co-treatment strategy.

Another prominent hit was DAPK3, detected as a soft-spot in half of the cell lines tested. DAPK3 is a member of the death-associated protein kinase family. It was found to have roles in induction of both apoptosis (44–46) and autophagic cell death (47). On the other hand, DAPK3 was also found to be important for tumor survival (48–50), in contrast to the well-known tumor suppressive properties of the DAPK family and their ability to promote cell death. Our results suggest that DAPK3 acts as a pro-survival protein in melanoma and therefore is a promising, relatively novel drug target. As a kinase, it is highly druggable, and several inhibitors have already been developed, although they do not show specificity within the DAPK family (51). The development of a more specific DAPK3 inhibitor would be warranted, since loss of expression and/or down-regulation of DAPK1 (and DAPK2) leads to tumorigenesis, and a wide acting inhibitor could potentially have contradictory effects on tumor growth and survival.

Altogether, we suggest that our strategy to analyze the functional death signature of individual tumors has the potential to dramatically change the state-of-the-art of combinatorial drug therapy in precision cancer treatment, by reducing the number of persister cells surviving the initial treatment with minimal overlapping toxicities, thereby lessening the odds of developing drug resistance at later stages, and preventing tumor relapse. While we based our siRNA platform on V600E *BRAF* early passage melanoma cell lines in combination with vemurafenib, it can readily be adapted to any tumor from which primary cultures can be generated for running the siRNA platform together with standard targeted drug treatment. Furthermore, expansion of the screen to include proteins from additional death modules, such as ferroptosis and parthanatos (52), will further increase our understanding of the intact death pathways that can be targeted to achieve maximal tumor killing. Currently, we employed clinically approved drugs to target soft-spots; however, with additional drugs in the pipeline, as well as RNAi protocols expected to be available for clinical use in the near future, more soft-spots can be harnessed for personalized, combinatorial treatment of melanoma and other cancer.

## Acknowledgments

We would like to thank, Aya Shkedy, Lital Povodovski, Rivi Halimi, Maya David Teitelbaum, Liat Hammer, Yuval Gilad, Gal Chaim Nuta, and Avital Hay-Koren for providing significant assistance and crucial advice with experimental procedures and writing of this paper.

## Notes

The authors declare no potential conflicts of interest

Funding: This work was supported by a grant from the Israel Science Foundation, Grant No. 679/17 to AK.

### Competing Interest Statement

The authors have declared no competing interest.

