## Supplemental Tables and Figures for "A functional pre-screening platform for identifying points of vulnerability in the cell death map of human melanoma tumors"

**Dekel-Bird, et al Supplementary Data**

**Supplementary Table S1**

**Supplementary Table S2 (legend, see excel file for table)**

**Supplementary Table S3**

**Supplementary Figure S1**

**Supplementary Figure S2**

**Supplementary Figure S3**

**Supplementary Figure S4**

**Supplementary Figure S5**

**Supplementary Table S1. Origin of 12 early passage melanoma cell lines**

| Line | Date of Surgery | Origin |
|---|---|---|
| 13 | 03/09/2006 | Lymph node |
| 14 | 19/09/2006 | Lymph node |
| 19 | 10/07/2007 | Subcutaneous |
| 29 | 29/08/2007 | Subcutaneous |
| 30 | 14/10/2007 | Lymph node |
| 31 | 11/12/2007 | Subcutaneous |
| 34 | 07/11/2007 | Lymph node |
| 39 | 20/12/2007 | Lymph node |
| 40 | 21/12/2007 | Subcutaneous |
| 41 | 01/04/2008 | Subcutaneous |
| 43 | 22/01/2008 | Subcutaneous |
| 50 | 18/09/2008 | Subcutaneous |

**Supplementary Table S2 Output of the siRNA viability screen (Excel file)**

* The ratio of the ln average of the treated divided by the ln average of the untreated value,

** *p*-values, comparing the change in viability of the siRNA KD to the si-control by a Dunnet test.

*** the change in viability of treated KD in comparison to the untreated KD of the same siRNA, by student T-test.

**Supplementary Table S3. List of hits in each cell line, classified according to modules in the PCD map**

| Cell Line | # Hits | Autophagy | Intrinsic apoptosis | Extrinsic apoptosis | Programmed necrosis | Other[a] |
|---|---|---|---|---|---|---|
| 13 | 2 | | BMF[b] | | | DAPK3 |
| 14 | 3 | ATG3 | MCL1, SURVIVIN | | | |
| 19 | 9 | ATG4B, ULK1 | BCLW, BCLXL, MCL1, CASP3, XIAP | **TRAF2** | **TRAF2** | DAPK3 |
| 29 | 1 | | BCLXL | | | |
| 30 | 11 | ATG3, ATG9A, VPS15, FIP200, UVRAG | BCLXL, MCL1, NOXA, PUMA | TRAILR2 | | DAPK3 |
| 31 | 2 | ULK2 | | **CFLIP** | **CFLIP** | |
| 34 | 9 | ATG10, ATG12, ATG5, FIP200, GABARAP | BIM, PUMA | **CFLIP** | **CFLIP** | DAPK3 |
| 39 | 6 | ATG5, ATG9A, VPS15 | MCL1, BNIP3L | FADD | | |
| 40 | 15 | AMBRA1, ATG5, FIP200, ULK1, VPS15, RHEB, GABARAPL1, WIPI2 | CASP6, PARP-1, PIDD, PUMA, XIAP | | RIPK1 | DAPK3 |
| 41 | 0 | | | | | |
| 43 | 14 | ATG3, GABARAP, GABARAPL1, MTOR | BAK, $BCLX_L$, CIAP1, NOXA, PUMA, XIAP | **TRAF2**, TRAILR1, TRAILR2 | **TRAF2** | DAPK3 |
| 50 | 2 | ATG16 | MCL1 | | | |

[a] Refers to members of the DAPK family, multi-functional regulators of various death pathways.

[b] Positive hits (soft-spots) are marked in black and negative hits in red. Dual function proteins participating in more than one module are in bold.

### Supplementary Figure S1

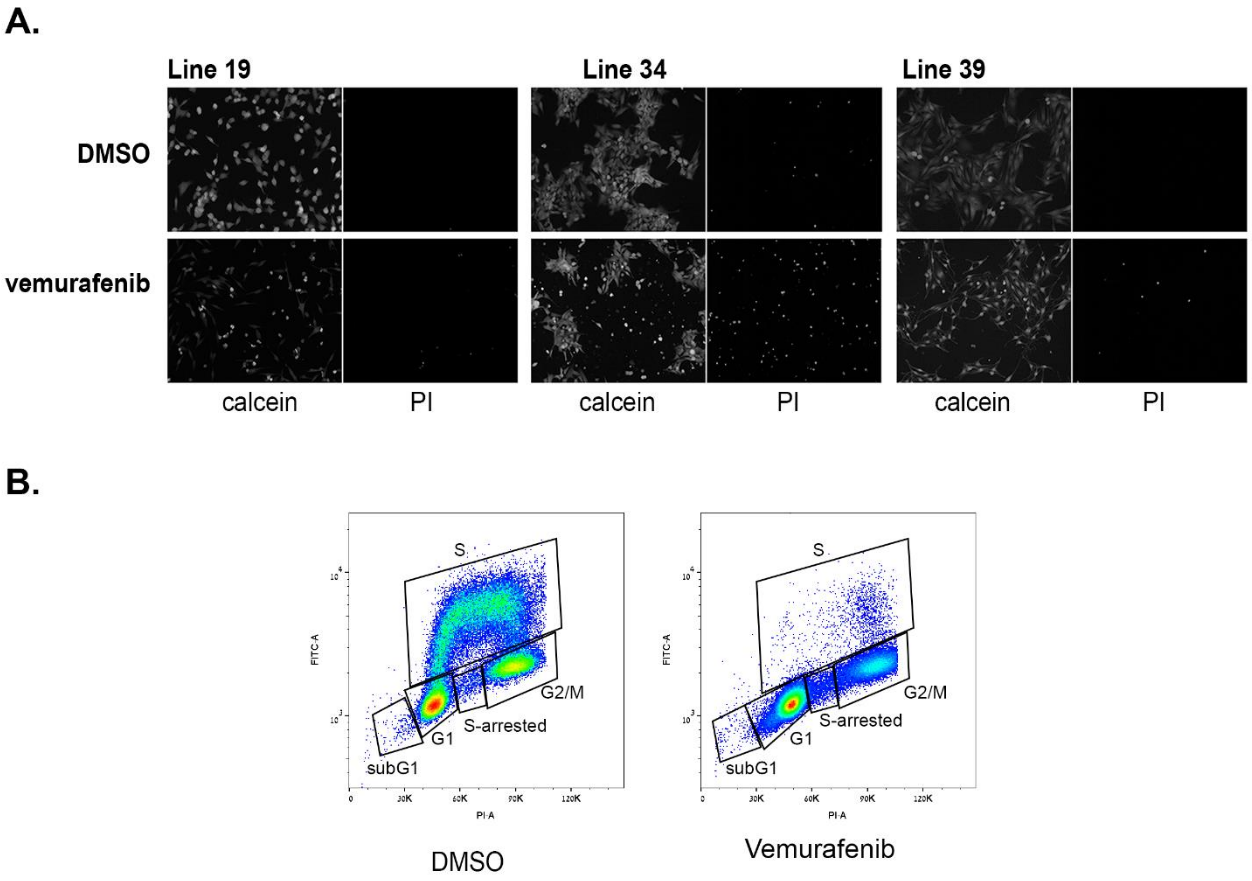


**Supplementary Figure S1. Patient-derived melanoma cell lines show heterogeneous responses to vemurafenib treatment. A.** Representative phenotypic response to vemurafenib of cell lines 19, 34 and 39 from Fig. 1. Left, single channel for the calcein AM (green) and right, single channel for PI (red), shown in greyscale. **B.** Cell cycle distribution of line 39 treated with 5μM vemurafenib for 24h stained with PI and anti-BrdU antibody. From each sample 50,000 cells were recorded. Shown is a representative experiment of three independent repetitions. The X-axis corresponds to PI staining and the Y-axis to anti-BrdU staining.

### Supplementary Figure S2

**A.**

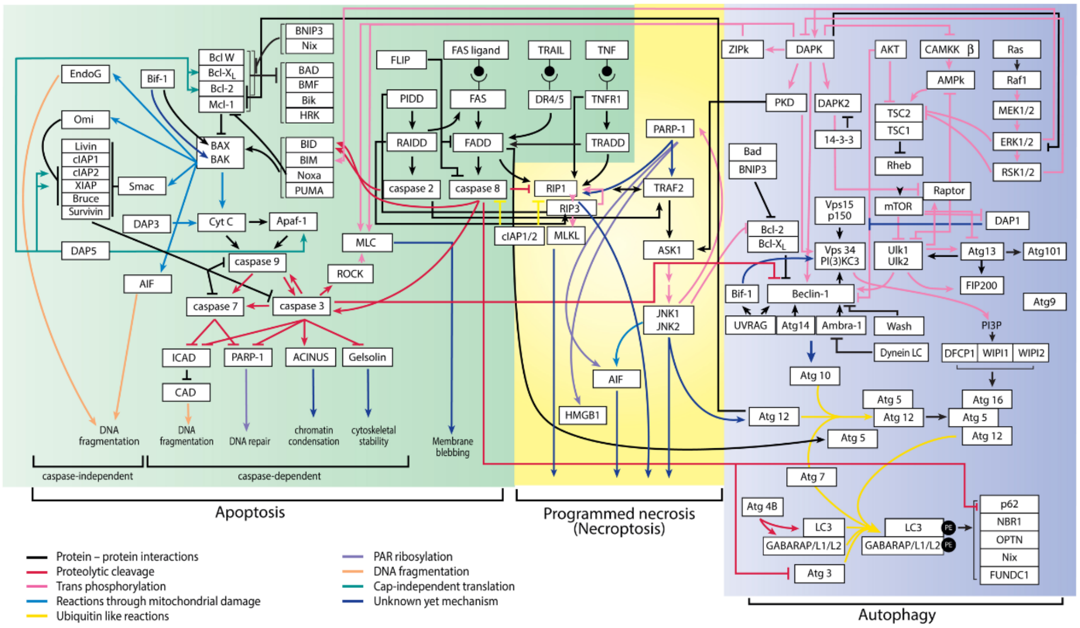


**B.**

| | 1 | 2 | 3 | 4 | 5 | 6 | 7 | 8 | 9 | 10 | 11 | 12 |
|---|---|---|---|---|---|---|---|---|---|---|---|---|
| A | Un-Tftd | Empty | Empty | Empty | Empty | Empty | Empty | Empty | Empty | Empty | Empty | Mock |
| B | Un-Tftd | Empty | MAP1LC3B | BAD | Empty | BNIP3 | NT5 | RISCfree | TRAF2 | MCL1 | Empty | Mock |
| C | Un-Tftd | BCL2 | CASP7 | DAPK2 | CASP8 | RIPK3 | BIM | ATG13 | ULK1 | VPS34 | NT2 | Mock |
| D | Un-Tftd | RISCfree | NT5 | RISCfree | FIP200 | NT2 | DAPK1 | ATG4A | BAX | DAPK3 | CASP9 | Mock |
| E | Un-Tftd | XIAP | BIK | ATG9A | BRUCE | APAF1 | GABARAPL1 | ATG7 | NT5 | PUMA | CIAP1 | Mock |
| F | Un-Tftd | AIF | TRAILR1 | RIPK1 | RHEB | ULK2 | PIDD1 | NT pool 2 | TRAILR3 | OMI | CIAP2 | Mock |
| G | Un-Tftd | Empty | CASP2 | NT2 | SMAC | WIPI2 | ATG12 | MLKL | CASP6 | BCL2A1 | Empty | Mock |
| H | Un-Tftd | Empty | Empty | Empty | Empty | Empty | Empty | Empty | Empty | Empty | Empty | Mock |

| | 1 | 2 | 3 | 4 | 5 | 6 | 7 | 8 | 9 | 10 | 11 | 12 |
|---|---|---|---|---|---|---|---|---|---|---|---|---|
| A | Empty | Empty | Empty | Empty | Empty | Empty | Empty | Empty | Empty | Empty | Empty | Empty |
| B | Un-Tftd | Empty | RAIDD | ATG3 | Empty | BCL2XL | NT5 | RISCfree | ATG14 | FADD | Empty | Mock |
| C | Un-Tftd | BID | ULK1 | MAP1LC3B | AMBRA1 | BECN1 | MTOR | ATG4B | RAPTOR | LIVIN | NT2 | Mock |
| D | Un-Tftd | RISCfree | NT5 | RISCfree | PARP1 | NT2 | BNIP3L | CASP10 | BCLW | ATG10 | TSC1 | Mock |
| E | Un-Tftd | CFLIP | WIPI1 | ATG5 | RIPK1 | TRAILR2 | SURVIVIN | ATG101 | NT5 | ATG12 | Empty | Mock |
| F | Un-Tftd | RIPK3 | TSC2 | VPS15 | DAP1 | CASP3 | BAK1 | NT pool 2 | UVRAG | GABARAP | GABARAPL2 | Mock |
| G | Un-Tftd | Empty | BMF | NT2 | NOXA | CASP8 | ATG16 | TRAILR4 | MCL1 | CIAP1 | Empty | Mock |
| H | Empty | Empty | Empty | Empty | Empty | Empty | Empty | Empty | Empty | Empty | Empty | Empty |

**Supplementary Figure S2. The Programmed Cell Death siRNA library A.** A programmed cell death (PCD) map delineating the landscape of protein-protein interactions within the three major modules of PCD, autophagy (highlighted in blue), programmed necrosis (highlighted in yellow) and apoptosis (highlighted in green), based on hundreds of curated publications. The edges possess directionality indicating either activation or inhibition of the target protein and are color coded as indicated. The map was initially published in Bialik S, Zalckvar E, Ber Y, Rubinstein AD and Kimchi A. Systems biology analysis of programmed cell death. Trends Biochem Sci 2010;35:556-564, and updated herein. **B.** Scheme of the siRNA coated plates used for identifying positive and negative hits in each cell line. The two 96-well plates contain siRNA pools targeting 81 PCD genes as well as si-controls (in red) distributed randomly on the custom designed plates (Dharmacon). Un-Tftd, untransfected cells; mock, transfected without siRNA added; unmarked wells were left empty.

### Supplementary Figure S3

**A.**

**Cell line 19**

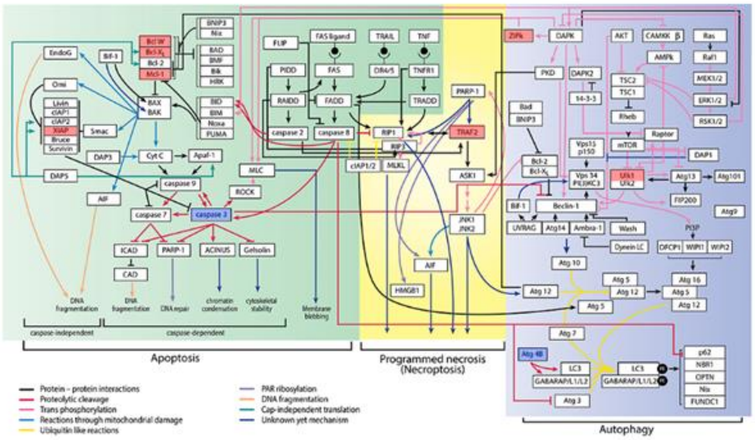


**B.**

**Cell line 34**

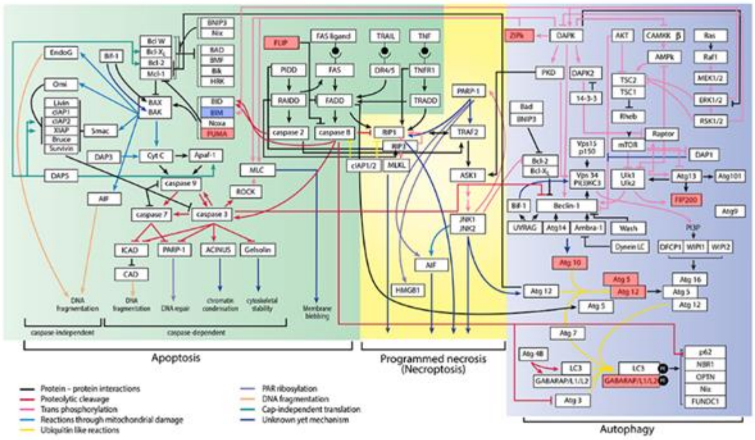


**C.**

**Cell line 39**

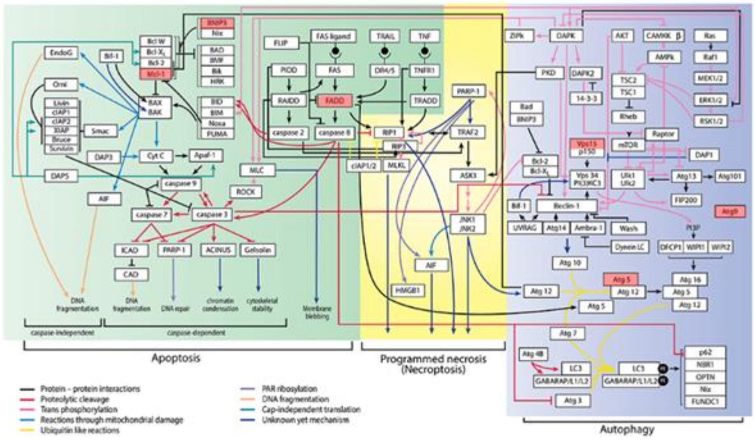


**Supplementary Figure S3. Cell death signature of melanoma.** Superimposing functional hits on the PCD map in cell lines 19 (**A**), 34 (**B**) and 39 (**C**). Positive hits (soft-spots) are marked in red and negative hits are marked in blue. The superimposition of the hits suggests open death pathways: for cell line 34 there is a cluster of positive autophagic hits suggesting blocking autophagy may reduce cell viability. In cell lines 39 and 19, the anti-apoptotic proteins from the BCL2 family seem to be the soft-spots.

### Supplementary Figure S4

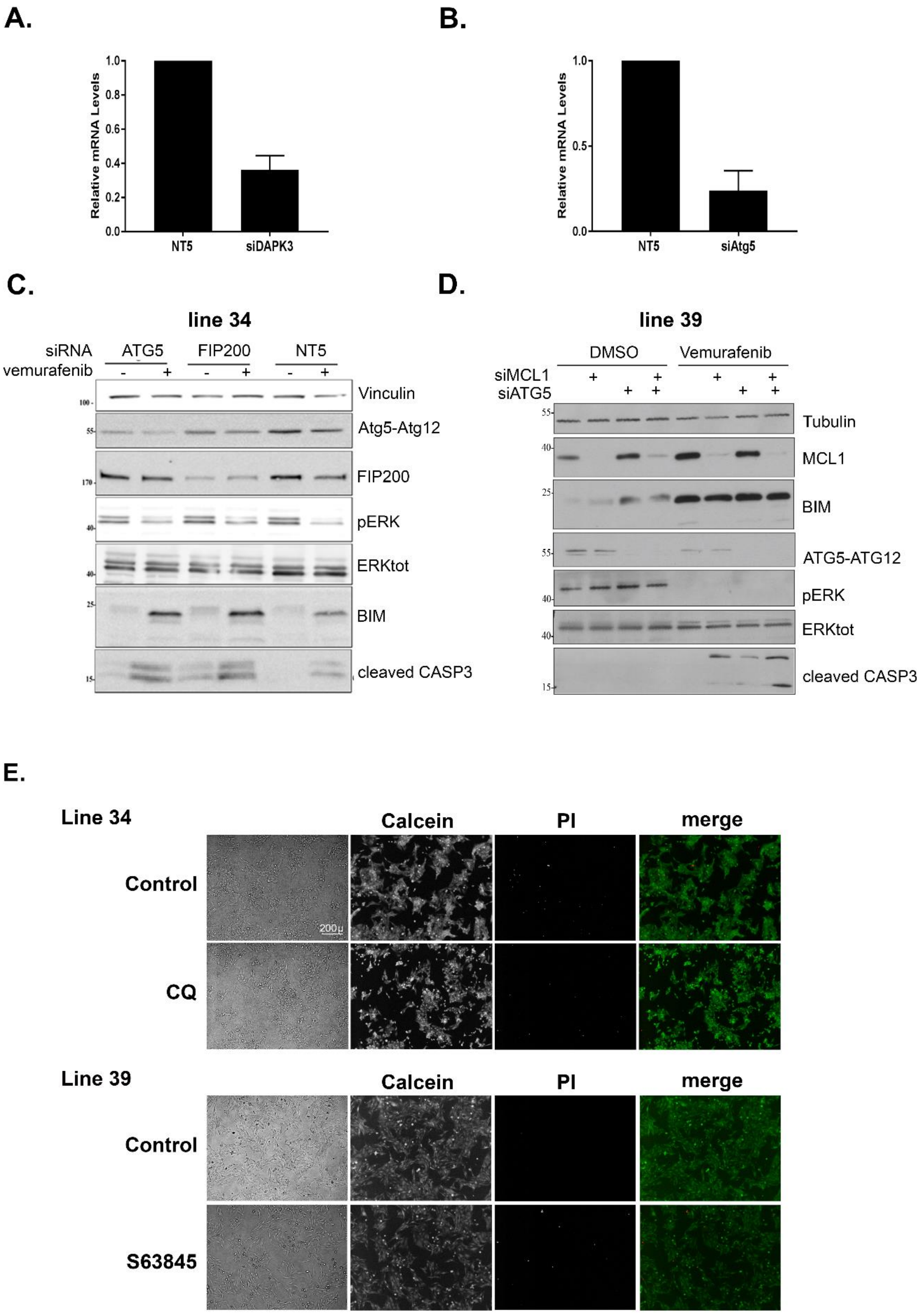

**Supplementary Figure S4. Validating points of vulnerability in the programmed cell death map in each tumor. A., B.** Quantitative RT-PCR analysis for DAPK3 mRNA levels (A) or ATG5 mRNA levels (B) in cell line 34 transfected with siRNA to DAPK3, ATG5 or non-targeting control siRNA (NT5). **C.** Western blot of line 34 transfected with siRNA targeting ATG5 or FIP200, either untreated or treated with 10µM vemurafenib. NT5 siRNA was used as control. **D.** Western blot of line 39 transfected with siRNA targeting ATG5 and MCL1, either untreated or treated with 10µM vemurafenib. RISCfree siRNA was used as control. In both blots, ATG12-ATG5 conjugate of the expected size is detected with anti-ATG12 antibodies. **E.** Lines 34 and 39 were treated for 24h with either 20µM chloroquine (CQ) or 0.5µM S63845, respectively, and then stained with calcein AM (green, stains live cells) and propidium iodide (PI) (red, stains cells with permeabilized membranes).

#### Supplementary Figure S5

A.

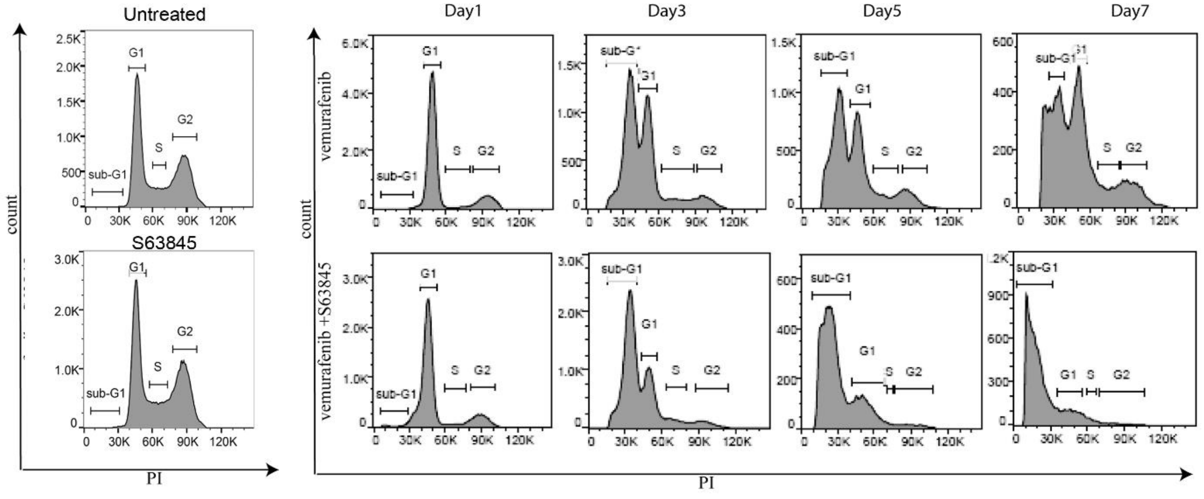


B.

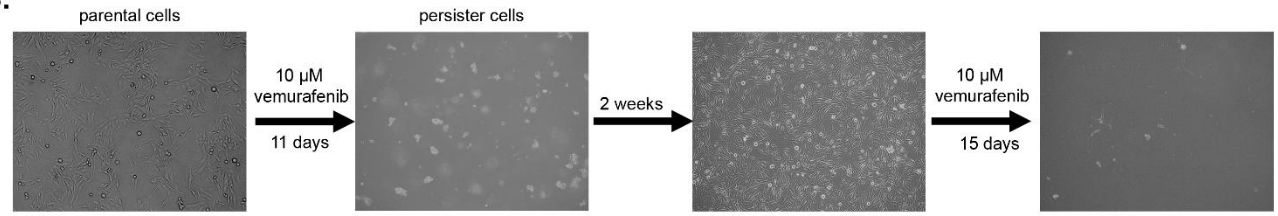


**Supplementary Figure S5. Targeting MCL1 during long term vemurafenib treatment of melanoma cell lines reduces viability of drug tolerant cells and the number of drug resistant foci. A.** Cell cycle distribution of cells from line 39 that were either untreated, or treated with 2µM vemurafenib, 0.5µM S63845 or both for 1, 3, 5 and 7 days. The Y-axis is cell number, the X-axis is PI staining. From each sample, 50,000 cells were acquired in 3 replicates. Percentage of the sub-G1 population is indicated. * $p<0.05$, ** $p<0.01$, combination vs vemurafenib alone. **B**. Representative images of response of parental and persister cells to long term vemurafenib treatment. Parental cell line 39 ($6x10^6$ cells per 15cm plates) was treated with 10µM vemurafenib for 11 days, resulting in killing of most of the cells. Upon drug removal, the remaining persister cells resumed cell growth, reaching confluency within two weeks. These cells were treated again with 10µM vemurafenib ($6x10^6$ cells per 15cm plates) for 15 days.
